# Haplotype-based fine-mapping of variant associations for complex traits

**DOI:** 10.64898/2026.10.19.760226

**Authors:** Arya Massarat, Michael Lamkin, Tara Mirmira, Utkarsh Jain, Jonathan Margoliash, Yang Li, Melissa Gymrek

## Abstract

Genome-wide association studies (GWAS) have identified tens of thousands of associations between genomic regions and complex traits in humans, but identifying the causal variants underlying these associations is challenging. Statistical fine-mapping techniques attempt to quantify the probability of causality of individual variants while accounting for the correlation induced by linkage disequilibrium (LD) between nearby variants. Existing fine-mapping solutions are limited to considering additive and independent contributions of individual variants, and also assume the causal variant is included in the analysis. Here, we introduce a novel fine-mapping method, happler, which models the effects of variant *haplotypes* (sets of alleles of two or more variants in *cis*) in addition to individual variants. By identifying haplotype proxies for causal variants, happler can infer additional effects including those driven by combinations of variants in *cis* and ungenotyped causal variants, such as complex structural variants, which are in some cases better tagged by one or more haplotypes than by individual SNPs. Simulation experiments show that whereas happler can correctly identify causal haplotype effects, traditional methods based on individual variants can produce inconclusive results in such cases. We applied happler to identify 32 haplotypes associated with gene expression using 359 individuals from the Geuvadis dataset. Of these, five are confidently fine-mapped, with stronger posterior inclusion probabilities than any individual variant, in an external cohort from the Genotype Tissue Expression (GTEx) project. Finally, we used the AoU dataset to demonstrate the utility of haplotypes for tagging rare causal variants that are absent from the input dataset.

Happler is available at: https://github.com/gymrek-lab/happler

## Introduction

Genome-wide association studies (GWAS) have identified tens of thousands of genomic loci associated with a range of traits in humans^1^. However, identifying the underlying causal variant(s) driving association signals is challenging. A typical GWAS signal may harbor hundreds of variants in strong linkage disequilibrium (LD) with each other, most of which are not causal for the trait. A variety of methods have been developed to perform statistical fine-mapping, with the goal of inferring the candidate causal variants at a particular GWAS locus, while accounting for LD^2^. However, existing methods face several important limitations: first, they assume the true causal variants exist within the set of candidate variants considered, which most often consists of common single nucleotide polymorphisms (SNPs) at a locus. Yet increasing evidence suggests that many of the strongest GWAS signals are driven by rare or complex variants that are in many cases not directly genotyped and may be in only modest LD with surrounding SNPs^3–5^. Second, they typically model independent contributions of individual variants rather than combinations of variants, which may either causally drive a signal or tag ungenotyped causal variants. Thus, existing fine-mapping solutions will not produce reliable results for such cases driven by ungenotyped variants or non-additive effects.

An alternative approach is to consider haplotypes, which we define here as sets of alleles in *cis*. Haplotypes can capture signals due to combinations of variants acting in *cis*^6,7^, and are often better able than individual SNPs to tag ungenotyped variants such as rare variants^8^, repeats^3^, or structural variants^9^. Haplotype tests^10–13^ were previously widely used when genotype data was limited to common SNPs and efficient phasing and imputation algorithms were not available, and thus the majority of variants were not directly tested. They declined in popularity after more complete reference haplotype panels of genetic variation based on next-generation sequencing became available, enabling accurate imputation of most common variants.

Even with the advent of large reference panels or biobank-scale whole genome sequencing, many effects including those driven by rare or complex variants or variant interactions are still not directly captured. Newer haplotype-based methods have been designed to further close this gap. For example, Howey and Cordell proposed using haplotypes composed of two SNPs for association testing, chosen based on their pairwise correlation, and used simulations to show that their approach is well-powered to capture many signals driven by ungenotyped variants^14^. Zheng *et al*. demonstrated that performing finemapping on haplotypes inferred from an external reference panel can improve effect size estimation at low sample sizes or when signals are driven by rare variants^15^. Bkhetan *et al*. constructed haplotypes based on LD between phased SNPs and recovered known signals that could not be detected by single SNPs^16^. Zhang et al used ARGs (ancestral recombination graphs) to infer the associations of unobserved variants and found that ARG-derived variants, which in most cases capture individual rare variants, can reveal additional associations missed by single variant approaches^17^. Although these methods use different strategies to define haplotypes, they all construct haplotypes based on LD information alone. While this has the benefit of obtaining phenotype-agnostic sets of haplotypes, it limits the portability of those haplotypes across training datasets that might have different LD structures. Further, it may not be the optimal strategy to identify haplotypes most relevant to specific phenotypes.

Here, we present happler, a new fine-mapping method which considers effects of haplotypes in addition to individual variants. Happler takes as input individual-level phenotypes and phased genotypes. Unlike previous approaches, it directly uses phenotype data to construct candidate trait-associated haplotypes at each region of interest. Candidate haplotypes are then used in addition to individual variants as input to an existing fine-mapping method to obtain a probability of causality for each variant and haplotype considered. We benchmarked happler using simulated data from haptools^18^, our open-source toolkit for simulation of SNP and haplotype effects. We demonstrate that happler remains sensitive across a variety of simulated effect sizes and allele configurations, whereas existing fine-mapping methods based only on individual variants can often produce inconclusive results, especially when the causal haplotype or variant is untyped. Finally, we apply happler to fine-map expression quantitative trait loci (eQTLs) for lymphoblastoid cell lines (LCLs) using the Geuvadis^19^ (discovery) and Genotype Tissue Expression Project^20^ (GTEx; replication) cohorts, and to GWAS loci for platelet count in the *All of Us* Research Program^21^. We identify specific signals at which haplotypes better explain variation in the trait than individual variants, suggesting that signals in those regions may instead be driven by complex variants, *cis* interaction effects, and rare variants not typically considered in GWAS and downstream analyses.

## Methods

### The happler method

We introduce a fine-mapping framework that takes as input a phased set of individual-level genotypes from a VCF or PGEN file of biallelic variants and phenotype values for a trait of interest and outputs posterior inclusion probabilities (PIPs, which quantify the probability of being causal) for each variant and a set of candidate haplotypes identified using the strategies described below (**Fig. 1A-B**). Happler consists of two stages: (i) haplotype construction, performed in a training cohort, followed by (ii) fine-mapping, performed in a validation cohort, in which haplotypes identified by the first stage are treated as additional variables input an existing fine-mapping tool, such as SuSiE^22^ (**Fig. 1B**). Happler operates on one locus at a time, considering only those genotypes in the locus surrounding each target lead variant. Typically, between 500kb-1Mb is chosen as the size of the locus, consistent with previous literature^23–25^.

**Fig 1.**
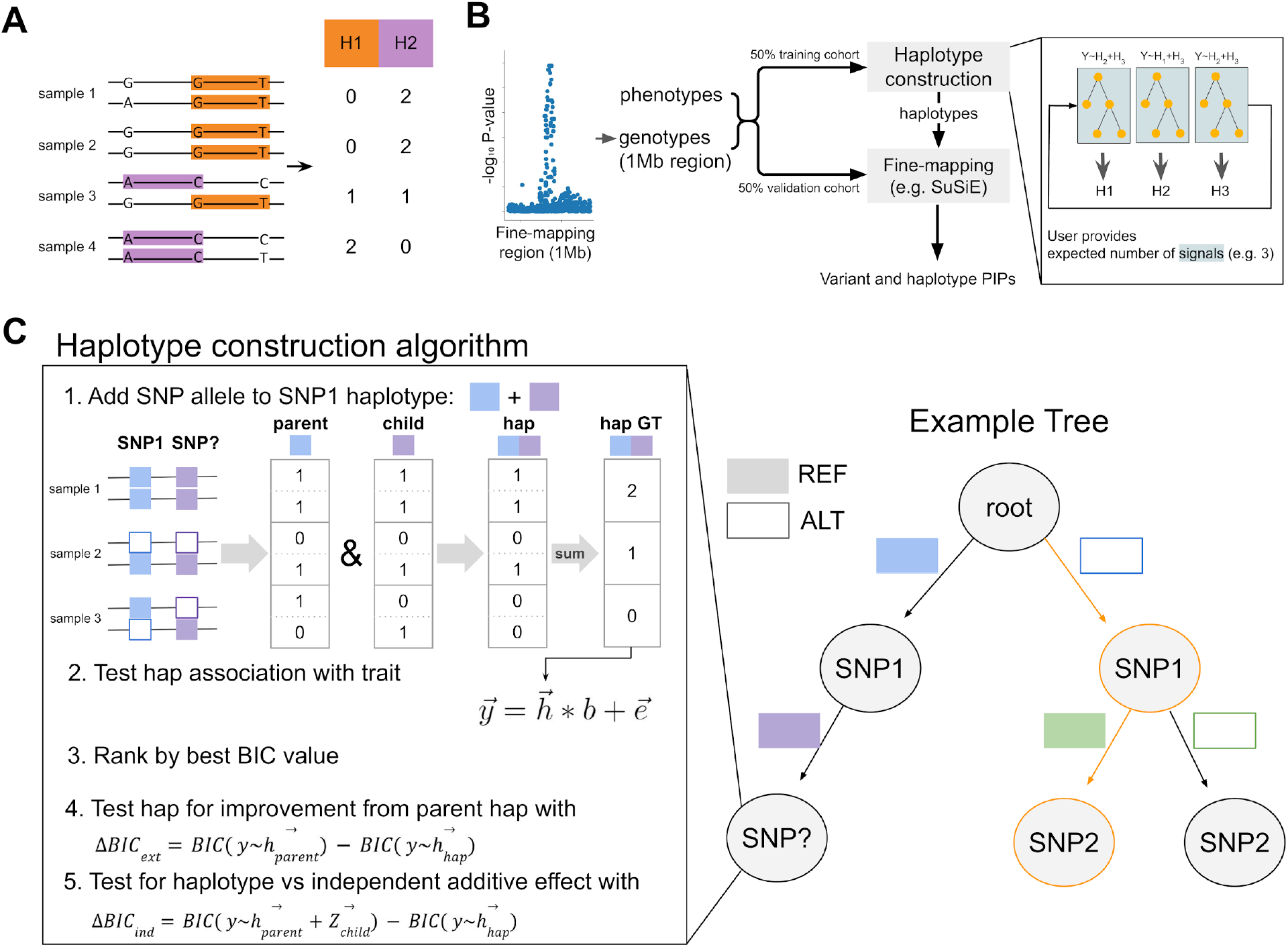
The happler method. A. Encoding haplotypes as individual-level genotypes. We define haplotypes as a set of two or more alleles in *cis*. These can be encoded as psuedo-genotypes by counting the number of times all alleles appear in *cis* across both chromosome copies. **B. Overview of the happler pipeline**. Happler consists of two stages performed in separate training and validation cohorts: *haplotype construction*, which is performed in the training cohort, identifies candidate trait-associated haplotypes, which are used alongside the input genotypes for individual variants as input to the *fine-mapping* stage, which uses the validation cohort. Inset: The method iteratively builds a tree for a user-specified maximum number of independent signals using residuals from regressing out haplotypes from other built trees as phenotypes. It repeats this process for a predefined number of iterations. **C. Overview of haplotype construction**. Within each iteration, the method builds a tree, where nodes of the tree are variants and branches are alleles. A path from the root of the tree to a leaf node (orange) represents a haplotype that would appear in the output of the method. At each node, we test whether to extend the current haplotype by each variant. The best variant is added to the tree if it is determined to improve the effect of the haplotype (based on Δ*BIC*_*ext*_) and if it is not determined to be explained by an independent additive signal (based on Δ*BIC*_*ind*_).

#### Haplotype construction

At each locus, happler uses phased input genotypes to identify a set of trait-associated *haplotypes*, where each haplotype is composed of a set of alleles at one or more variants that occur in *cis* (**Fig. 1A**). The alleles are greedily chosen to maximize the strength of the association between the haplotype and the target trait. To identify candidate haplotypes, we construct a binary tree composed of linear regression models (**Fig. 1B-C**). Each node in the tree represents a variant with branches of a node representing the variant’s alleles: REF or ALT. A path from the root of the tree to a target node *V* passes through alleles of one or more variants, which defines the *haplotype* ending at node *V*. Note, a haplotype consisting of only a single node, arising from a tree with only one non-root node, is equivalent to an individual variant.

The algorithm begins by choosing the variant with the strongest trait association to place at the root node of the tree. Association tests are performed using a simple linear model for each variant 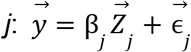, where 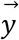 is a vector of individual-level phenotypes, 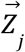 is a vector of individual-level normalized genotypes for variant *j*, β_*j*_ is the unknown effect size of variant *j*, and 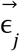 is a vector of noise terms. Phenotypes are assumed to already be adjusted for relevant covariates. Genotypes are Z-normalized to have mean 0 and variance 1. The strongest trait association is determined based on the Bayesian Information Criterion (BIC).

Non-root nodes are constructed according to a different strategy. For each node *V*_*parent*_ in the tree, we evaluate whether extending the haplotype of that node by each possible variant allele not already in the haplotype will improve the model fit. For each possible haplotype extension, we construct a *haplotype matrix* encoding whether each individual has 0, 1, or 2 copies of the candidate haplotype (**Fig. 1C**). The haplotype matrix is analogous to the genotype matrix that is traditionally used for association testing in GWAS and fine-mapping methods. To determine the best allele for the next node *V*_*child*_ of the tree, each new haplotype is tested for an association with the phenotype using the model above, except the 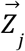 term encodes counts from the haplotype matrix rather than variant-level genotypes. Alleles that would reduce the MAF of the haplotype below a user-defined threshold can optionally be excluded from consideration. The allele from the haplotype with the best p-value is chosen as the candidate allele for *V*_*child*_.

#### Termination criteria

Before extending the haplotype by the best allele identified above, we perform two additional tests. Imagine we are considering whether to add a new allele at a node *V*_*child*_ in the tree. The standardized genotypes of the new allele are given by 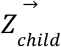, and those of the haplotype at the parent node are given by 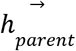. Then 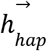 represents the haplotype at *V*_*child*_, which is the extension (logical AND) of 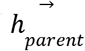 and 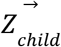. First, to determine whether the addition of the allele to the haplotype improves the model fit compared to the parent haplotype, we compute the delta BIC of the extension, 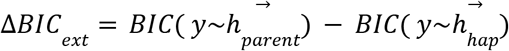 (**Fig. 1C**). We also compute a separate delta BIC to determine whether the candidate extended haplotype outperforms a model in which the parent haplotype and candidate node have independent effects, 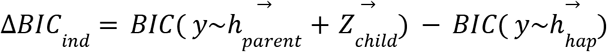. If one or both metrics do not exceed a specified threshold, then tree building is terminated. Default threshold values were identified based on simulation analyses described below.

#### Handling multiple independent signals

Many trait-associated regions have been shown to be driven by multiple independent signals^26^. To allow for this case, we perform an iterative procedure, similar to that used in Algorithm 1 of the SuSiE method^22^, in which we build multiple trees, each corresponding to a potential independent signal, where the user specifies a maximum number of causal signals (*L*) for the locus. For *i* in 1 … *L*, we build a separate haplotype tree, using phenotype values for which we have regressed out all candidate haplotypes already discovered by other trees (**Fig. 1B**). We repeat this process for a user-specified maximum number of iterations (default 3) and return the union of haplotypes from each tree.

The final set of haplotypes output by each tree is filtered to exclude: those with association p-values below a user-specified threshold (default p=5*10-8), and those consisting of a single variant, since all individual-level variants are in any case provided as input to the fine-mapping stage.

#### Fine-mapping to compute haplotype and variant level PIPs

In the final step of the method, we encode the haplotypes at the leaves of the tree in a haplotype matrix. The resulting matrix is used in addition to variant-level genotypes as input to an existing fine-mapping method to compute a PIP for each haplotype and each variant. Although in theory any fine-mapper can be used, in practice we use SuSiE^22^ over FINEMAP^24^ because it explores more causal configurations and has demonstrated improvements in speed.

#### Implementation

Happler is implemented as an open-source Python package. All file IO is powered by the haptools^18^ data library and all linear associations were performed with statsmodels.regression.linear_model.OLS^27^.

### Simulation analyses

To benchmark our method, we first use simulated phenotypes. To ensure our simulated data recapitulates realistic LD patterns, we constructed haplotypes from real genotypes in the 1000 Genomes Project Phase 3^28^ dataset distributed on the PLINK2^29^ website in the locus with coordinates chr19:45401409-46401409. Only bi-allelic SNPs with alternate allele frequencies between 0.05 and 0.95 were considered. We randomly selected pairs of alleles, each from a separate SNP, to generate haplotypes. The LD between the alleles of each haplotype was computed as the Pearson correlation coefficient. Then, the resulting haplotypes were filtered so that the final set has a roughly even distribution of LD values between the two alleles defining the haplotype. For this, we assigned haplotypes to 10 equally spaced bins between 0 and 1 based on the LD of their alleles and chose 1 haplotype at random from each bin for a total of 10 haplotypes.

We used haptools^18^ v0.6.2 to simulate phenotypes from these haplotypes according to the linear model 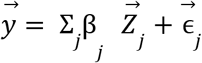 where 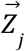 corresponds to the standardized counts of haplotype *j* for all individuals and the errors are sampled randomly from a normal distribution and assumed to be independent for each sample: 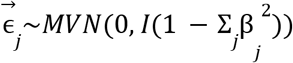. We repeated this process 10 times each for values of β_*j*_ of [0.05, 0.1, 0.15, 0.2, 0.25, 0.3]. We evaluated scenarios with between 1 and 4 total causal haplotypes and random sample subsets of [800, 1600, 2400, 3200]. For each setting, we repeated two rounds of simulation, using one dataset for haplotype construction and the other as the orthogonal replication dataset for fine-mapping. Happler was executed with at most L=3 trees and 3 iterations, and its haplotypes were constructed to have a minimum MAF of 0.05. We additionally filtered haplotypes not reaching P<5e-8.

We measured the performance of happler by computing the LD between each observed haplotype output by happler and its best match among the causal (simulated) haplotypes. Observed haplotypes were uniquely paired to causal haplotypes according to a linear assignment via scipy.optimize.linear_sum_assignment, which implements minimum weight matching in bipartite graphs. This assignment maximizes the total LD between the pairings while ensuring that a causal haplotype is never paired with two or more observed haplotypes. When happler reported more observed haplotypes than the number of true causal haplotypes, we reported the remaining haplotype that could not be paired separately. After haplotype construction, observed haplotypes as well as individual genotypes and phenotypes from the replication dataset were used as input to SuSiE v0.14.2 to perform fine-mapping under default settings with L=10.

### Parameter tuning and benchmarking

We extended our simulation framework to benchmark and tune thresholds (Δ*BIC*_*ext*_, Δ*BIC*_*ind*_) for the two model comparison steps used by happler at each node in the tree-building process. We randomly subsetted the 1000 Genomes data^30^ to four sets of 800, 1600, 2400, and 3200 samples. Causal haplotypes within each Geuvadis eQTL locus (described below) were randomly constructed from pairs of alleles using the process described above at the lowest sample size of 800. The resulting haplotypes were filtered to ensure an approximately even distribution of LD values between the two alleles defining the haplotype. For this, we assigned haplotypes to 10 equally spaced bins between 0.05 and 0.95 based on the LD of their alleles and chose 10 haplotypes at random from each bin for a total of 100 haplotypes. We used haptools to simulate phenotypes according to the model described above. Each haplotype was used to simulate phenotypes with effect sizes (β_*j*_) of [0.05, 0.1, 0.15, 0.2, 0.25, 0.3]. This process was repeated five times for each of the four sample subsets.

First, we sought to optimize and evaluate the BIC threshold used when deciding whether to extend a node of the tree. For each simulated phenotype, we computed the BIC value of the model with the haplotype *BIC*(*y*~*h*_*hap*_) and the BIC value of the model with only the first allele in the haplotype *BIC*(*y*~*h*_*parent*_). The alleles of each haplotype are ordered such that the allele with the best (lowest) BIC value based on individual association tests for each allele appears first. We also extended the haplotype by the next best node in the tree (so the haplotype now has three alleles) and computed the BIC a third time: *BIC*(*y*~*h*_*extension*_). For each simulated phenotype, we labeled the Δ*BIC* of *BIC*(*y*~*h*_*parent*_) − *BIC*(*y*~*h*_*hap*_) as a “positive” (since *h*_*hap*_ is the true causal haplotype) and the Δ*BIC* of *BIC*(*y*~*h*_*extension*_) − *BIC*(*y*~*h*_*hap*_) as a “negative” (since extending would result in an incorrect haplotype; **Supplementary Fig. 1**). We evaluated the ability of the Δ*BIC*_*ext*_ score to classify between positives and negatives by computing the performance metrics AUROC and average precision.

We used a similar process to evaluate the performance of the test for an independent association. For this, the positives corresponded with simulations in which the haplotype was simulated as causal *y*~*h*_*hap*_, while the negatives represented simulations in which the parent of the haplotype and the haplotype’s last allele were additively causal as in *y*~*h*_*parent*_ + *Z*_*child*_. For both cases, we used the Δ*BIC*_*ind*_ of those two models, *BIC*(*y*~*h*_*parent*_ + *Z*_*child*_) − *BIC*(*y*~*h*_*hap*_), as the scoring metric (**Supplementary Fig. 1**). We evaluated the ability of this score to classify between the positive and negative cases by computing the same performance metrics.

To choose appropriate thresholds for both Δ*BIC*_*ext*_ and Δ*BIC*_*ind*_, for each simulation setting we determined Δ*BIC*_*ext*_ or Δ*BIC* _*ind*_ thresholds that result in a false discovery rate (FDR) of at most 5%. We found optimal thresholds in each setting are relatively robust to changes in sample size, with Δ*BIC* _*ext*_ thresholds mostly below 18 and Δ*BIC*_*ind*_ all below 14. We therefore defined a default setting of Δ*BIC*_*ext*_ > 20 and Δ*BIC* _*ind*_ > 15. After executing happler, we also recompute a final value of Δ*BIC*_*ext*_ (Δ*BIC*_*ext, final*_) against the original phenotypes. This differs from the Δ*BIC*_*ext*_ values computed during each iteration, which are based on phenotypes that have been adjusted by candidate signals arising from other trees in the forest. Overall, we found that Δ*BIC*_*ext, final*_ of the output haplotype is a useful quality metric, with high Δ*BIC* _*ext, final*_ values indicating the highest confidence haplotypes most likely to replicate in orthogonal cohorts.

### Geuvadis data and preprocessing

Previously computed expression quantitative trait loci (eQTL) summary statistics based on 359 individuals of European ancestry were obtained from the Geuvadis gene expression project at https://ftp.ebi.ac.uk/biostudies/fire/E-GEUV-/001/E-GEUV-1/Files/E-GEUV-1/analysis_results/EUR373.gene.cis.FDR5.best.rs137.txt.gz^31^. We lifted over 2,440 eQTLs identified at FDR 5% corresponding to 12,606 unique genes from hg19 to hg38 using the UCSC liftover^32^ tool v1.3.1 with chain file https://hgdownload.soe.ucsc.edu/gbdb/hg19/liftOver/hg19ToHg38.over.chain.gz. Individual-level gene expression values were obtained from https://www.ebi.ac.uk/biostudies/files/E-GEUV-1/E-GEUV-1/analysis_results/GD660.GeneQuantRPKM.txt.gz^31^. Individual-level phased genotypes from the 1000 Genomes Project^33^ corresponding to these individuals were obtained from https://www.internationalgenome.org/data-portal/data-collection/30x-grch38. Prior to fine-mapping, we regressed out the following covariates from gene expression values: 36 PEER factors^34^, sex, and the first 5 genotype PCs, where the PCs were computed as part of a previous study. We fine-mapped 1 Mbp regions centered around each eQTL. For downstream processing, we used PLINK2 v2.00a5.10 to extract only biallelic SNPs with a minimum minor allele frequency (MAF) of 0.1. Happler was executed with at most L=3 trees and 3 iterations, and its haplotypes were constructed to have a minimum MAF of 0.1 and maximum p-value of 5e-6.

We computed the *R*^2^ of the model fit via statsmodels.regression.linear_model.OLSResults.rsquared^27^ for each candidate haplotype based on a linear model 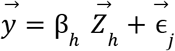 with standardized haplotype counts 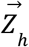 and phenotypes 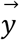 (**Supplementary Table 1**). For comparison we built a second model consisting of additive and independent contributions of the alleles defining each haplotype: 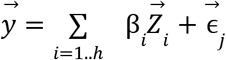 where 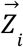 represents the genotypes for variants corresponding to the *i*th allele used to construct the haplotype and *h* is the total number of alleles in the haplotype (typically two, but can in some cases be more than two). The *R*^2^ of the model fit was computed in the same way as described above.

### GTEx data and preprocessing

We utilized available gene expression and genotype data from the Genotype-Tissue Expression (GTEx) Project^35^ v8 available through dbGaP accession number phs000424.v8.p2 to perform fine-mapping using haplotypes identified in the Geuvadis cohort. We focused on 131 samples with expression data available for EBV Transformed Lymphocytes. SNP Genotypes used in this study were from dbgap (file GTEx_Analysis_2017-06-05_v8_WholeGenomeSeq_866Indiv.vcf.gz), and filtered using PLINK2 with the following parameters: --hwe 0.001 and --geno 0. We then performed principal component analysis (PCA) of available SNP genotypes using PLINK2^29^ to infer the set of samples with predicted European ancestry. Based on visual inspection of the first two genotype PCs, we classified a sample as European if PC1>-0.014 and PC2 > 0.01 resulting in 109 European samples.

Inverse normal transformed expression values for each gene were obtained from the GTEx website (GTEx_Analysis_v8_eQTL_expression_matrices.tar). We computed PEER factors using PEER^34^ v1.0. For PEER factor computations, we further removed genes with TPM<1 in more than 90% of individuals. Prior to downstream analyses we regressed out the top 5 genotype principal components (PCs), sex, and 11 PEER factors. For each candidate haplotype-gene pair, we computed the *R*^2^ of the model fit using an additive model including both of the individual SNPs contributing to each haplotype as well as the haplotype itself. SuSiE v0.14.2 was used for the fine-mapping step with L=10.

### All of Us data and preprocessing

We applied happler to fine-map GWAS signals for platelet count identified in the *All of Us* Research Program^21^ v8 dataset (AoU). As described previously^36^, prior to downstream analysis, we removed samples whose value for sequencing metrics ins_del_ratio, snp_het_homvar_ratio, indel_het_homvar_ratio, ti_tv_ratio, or singleton metrics falls outside 8 standard deviations of the means across all samples. We also subsetted to only the samples that were included within AoU’s list of unrelated individuals and which had passed sex concordance checks performed by AoU. We restricted to samples which had self-identified race as White and had genetically inferred ancestry as ‘eur’ according to AoU’s ancestry prediction file. We focused on this cohort since it had the largest available sample size compared to other ancestry groups.

We then obtained data corresponding to concept ID 3024929 (“Platelets [#/volume] in Blood by Automated count”). Data points with units “thousand per microliter”, “per cubic millimeter”, or “/mm3” were kept. We additionally kept data points with “no value” or “No matching concept” listed as the unit since these followed a similar distribution to the retained measurements. After restricting to data points from individuals passing sample preprocessing above, we filtered values more than 5 standard deviations away from the mean. We then obtained the median phenotype value per year that measurements were available, then took the median of medians across all years. We recorded the year corresponding to the final value used as the age covariate for downstream association testing. After preprocessing, a total of 105,446 samples remained for analysis.

We applied fine-mapping to regions we previously identified in Mirmira *et al*.^*36*^ for platelet count in AoU. Before running the happler workflow, we regressed out all covariates (first 10 PCs, age, and sex at birth) from the normalized trait values. We ran happler with the phased short-read SNP/indel VCF files provided by AoU. Using PLINK2^29^, we removed multiallelic variants and variants with HWE p-value less than 1e-100. Happler was initially run with the following parameters: --hap-maf 0.0001 --max-signals 3 --max-iterations 3 --discard-multiallelic --remove-SNPs --indep-thresh 15 -t 20 --chunk-size 500 --out-thresh 5e-08. For subsequent analysis, to identify candidate causal variants, we ran SuSiE v0.14.2 with default parameters including all bi-allelic variants with MAF>0.0001 as input.

## Results

### Benchmarking happler using simulated data

We first evaluated happler under a range of simulation settings in which we simulated phenotypes assuming a single causal haplotype and varied the simulated effect size (range 0.05-0.3), sample size (range 800-3200), and LD between alleles (range 0-1) of each causal haplotype. In each case, samples were randomly split 50/50 into training (haplotype construction) and validation (fine-mapping) cohorts. We first measured the performance of the *haplotype construction step* of happler by computing the absolute value of the Pearson correlation between the best output (observed) haplotype and causal (simulated) haplotypes. As expected, the correlation with the causal haplotype tends to increase at higher effect sizes and larger sample sizes (**Fig 2A**).

**Fig 2.**
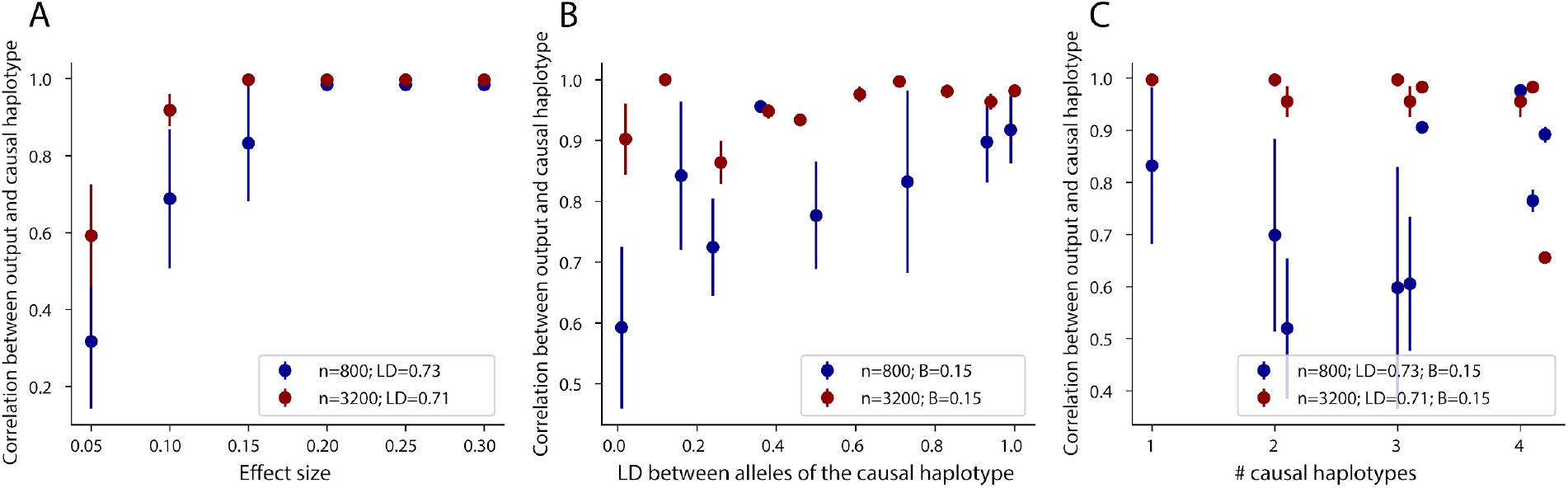
Benchmarking happler’s haplotype construction step using simulations. We simulated traits based on causal 2-SNP haplotypes with a range of effect sizes, sample sizes, and LD values between the two SNPs of each haplotype. We used happler to construct candidate haplotypes for each simulated trait and evaluated performance by computing the correlation between output haplotypes and simulated causal haplotypes. For **A-C**, results are shown for two sample sizes: 800 (dark blue) and 3200 (dark red). In all cases dots give the mean and error bars denote +/-1 standard error based on 10 replicates. In each of the 10 replicates the simulated causal haplotype is the same but the random seed is changed. **A. Correlation between the output and causal haplotype vs. effect size**. The x-axis gives the simulated effect size and the y-axis gives the correlation with the causal haplotype. The legend indicates the LD between the alleles of the haplotype for the two simulated sample sizes. **B. Correlation between the output and causal haplotype vs. the LD between the haplotype alleles**. The x-axis gives the LD between the two alleles and the y-axis gives the correlation of the output with the causal haplotype. For both sample sizes, results are shown for simulations with an effect size of 0.15. **C. Correlation between output and causal haplotypes in the case of multiple causal haplotypes**. We simulated between 1-4 causal haplotypes but performed haplotype construction allowing for a maximum of 3 signals. Output haplotypes were matched with simulated haplotypes using a procedure described in **Methods** and the plot shows the correlation with the matched causal haplotype for each output haplotype. For all panels, correlation is computed as the absolute value of the Pearson correlation coefficient.

We found that the correlation of the output haplotype with the causal haplotype tends to increase with the LD between alleles on the causal haplotype (**Fig. 2B**). In cases for which the alleles of the haplotype are in high LD, happler tends to output haplotypes containing only a single allele, which is already a strong proxy for the causal haplotype. Haplotypes for which the alleles are in modest LD showed lower correlation, in most cases because the termination tests (**Methods**) required to extend the haplotype to the second allele did not achieve the required statistical significance. As expected, this trend improves with increasing sample size (**Fig. 2B**) which results in improved power for both tests. Full simulation results are shown in **Supplementary Fig. 2**.

We next evaluated happler’s performance in cases of multiple, independent causal haplotypes and tested the effect of the input parameter *L* specifying the maximum number of signals to identify. We simulated between 1-4 causal haplotypes per locus while allowing happler to build *L*=3 trees. In cases where the number of causal haplotypes was less than or equal to *L*, haplotype construction identified haplotypes that were strongly correlated with each of the input haplotypes (**Fig. 2C**), indicating the iterative procedure is correctly identifying independent signals. As expected, when the number of trees built was higher than the number of simulated signals, not all output haplotypes were matched to a causal haplotype. Further, when simulating 4 causal haplotypes but setting L=3, only three matching haplotypes were identified.

We then evaluated the performance of the *fine-mapping step* of happler, which uses variant-level genotypes in addition to candidate haplotypes output during the haplotype construction step as input to an existing fine-mapping tool (in this case, SuSiE^22^). This step allows us to obtain PIPs for the haplotypes and assess whether the haplotypes are better candidates for the causal variant than any other individual variant (**Fig. 1B**). We first assessed the false positive rate of the fine-mapping step on the original (training) cohort vs. on an independent validation cohort. For this, we performed additional simulations in which the true causal variant is an individual SNP, rather than a haplotype. We then performed haplotype construction on the training cohort to obtain the best candidate haplotypes. Finally we evaluated whether the PIP of a simulated causal SNP would be affected by inclusion of a non-causal haplotype (**Supplementary Fig. 3**). The PIP remains unaffected in the validation cohort, whereas fine-mapping in the training cohort incorrectly assigns lower PIPs to the truly causal SNP, often incorrectly assigning the non-causal haplotype a high PIP. In both cohorts, the rate at which haplotypes were falsely identified as causal (at a PIP threshold of 0.5) is lower at higher delta BIC thresholds and is most attenuated when the fine-mapping step is performed in a separate validation cohort (**Supplementary Fig. 3**). Taken together, these results indicate that using an orthogonal validation cohort for fine-mapping is critical to avoid assigning non-causal haplotypes high PIPs due to over-fitting.

After establishing the need to perform fine-mapping in an independent cohort, we next assessed the performance of fine-mapping in this cohort using simulations described above in which a haplotype is the true causal variant. Haplotypes obtained from the discovery cohort in addition to individual variants were used as input to SuSiE for fine-mapping. We found that as expected the PIP of candidate haplotypes increases with effect size. Further, fine-mapping could better identify the haplotype as causal in cases where the LD between the haplotype’s alleles is modest (**Fig. 3A**), since in the case of high LD the individual alleles of the haplotype are in high LD with the haplotype itself, making it difficult for fine-mapping to distinguish between them. For comparison, we performed traditional fine-mapping using SuSiE with only individual variants as input. As expected, in cases of modest LD between the haplotype alleles, happler correctly identifies the candidate haplotype as causal with high posterior probability (**Fig. 3B**), whereas SuSiE based on variants alone is not able to identify a strong candidate causal variant (**Fig. 3C**).

**Fig 3.**
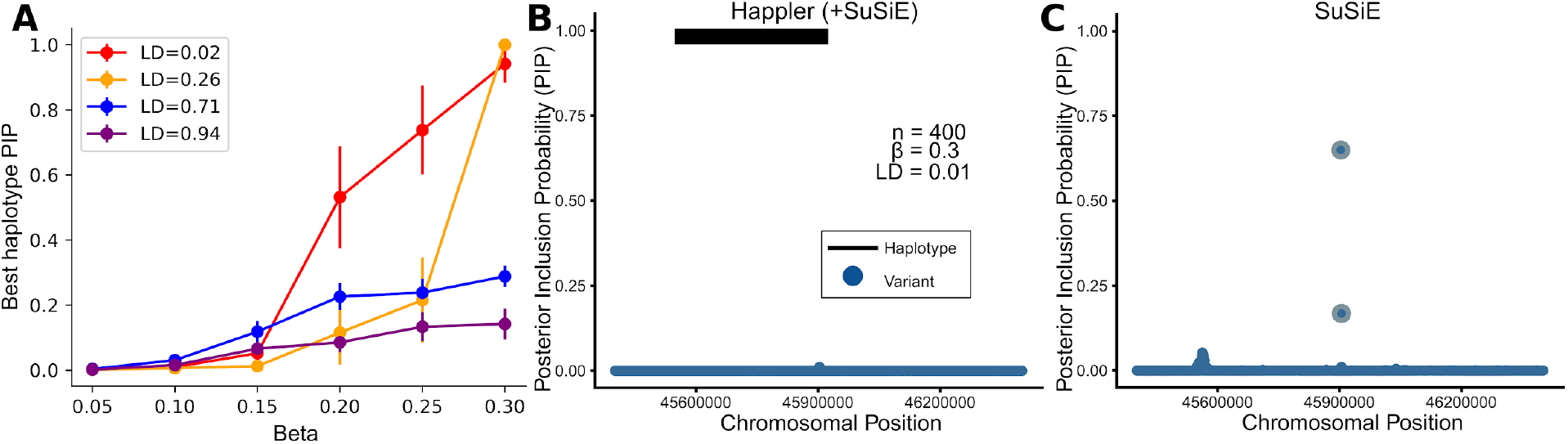
Benchmarking haplotype-based fine-mapping with simulations. Observed haplotypes identified during the haplotype construction step in the training dataset, plus individual variants and simulated phenotypes described above were input to SuSiE for fine-mapping in the validation dataset. Plots show the case of a single simulated causal haplotype. **A. Effect size vs. posterior inclusion probability (PIP)**. The x-axis represents the simulated effect size (Beta) and the y-axis represents the best PIP obtained for an observed haplotype. Dots give the mean across 10 simulations and error bars represent +/-1 standard error. Colors denote the LD between the two alleles of the simulated haplotype. Phenotypes were simulated for n=1600 samples. **B. Happler PIPs for all variants in an example region**. PIPs are shown for all individual variants (blue) and the observed haplotype (black bar spanning the location of the two alleles defining the haplotype) in an example fine-mapped region with simulation parameters LD between alleles=0.01, n=400, Effect size=0.3. **C. SuSiE PIPs for all variants in the same example region**. The plot is the same as in **B**, but based on running SuSiE with only variants (no haplotypes) as input. In this case SuSiE fails to identify a high confidence causal variant.

### Haplotype-based fine-mapping eQTLs in the Geuvadis cohort

We applied happler to fine-map eQTL signals identified in lymphoblastoid cell lines (LCLs) for 359 European samples for which RNA-sequencing is available from the Geuvadis^19^ cohort and genotypes are available from the 1000 Genomes Project^33^. We focused on 2,406 genes for which significant eQTLs were previously identified and performed fine-mapping in a 1Mb region centered on each lead SNP. Haplotype construction identified candidate haplotypes at a total of 32 eQTL loci under default settings (**Supplementary Table 1**). Of the 32 candidate haplotypes, 31 haplotypes consisted of two alleles, and a single haplotype contained 3 alleles.

Next, we evaluated candidate haplotypes in an independent cohort using available LCL expression data and matched genotypes available for 109 samples of predicted European ancestry from the GTEx Project^35^ (**Supplementary Table 1**). Of the 32 genes with candidate haplotypes, 30 were present and could be assessed in GTEx. Effect sizes of candidate haplotypes are highly correlated across datasets (Pearson *R*^2^=0.95; two-sided P=3.4e-15; n=31; **Supplementary Fig. 4a**). We observed that the majority of haplotypes with Δ*BIC*_*ext, final*_ > 25 exhibited a higher model R^2^ than their individual alleles in GTEx (14/19, 73.7%) (**Supplementary Fig. 4b**), indicating that the haplotype better explains the signal compared to its component alleles even in an independent cohort. Similarly, haplotype frequencies and the LD between alleles in each haplotype were highly consistent across datasets (**Supplementary Fig. 4c-d**).

We used the GTEx cohort to perform fine-mapping for each haplotype identified in Geuvadis, using both the haplotypes and the individual variants as input for each gene (**Fig. 4A**). Of the 14 total haplotypes identified above exhibiting a higher model R^2^ than their component alleles, five haplotypes obtained PIPs above 0.9 in GTEx (**Supplementary Table 1)**. All other tested haplotypes did not achieve PIPs above this threshold. These haplotypes were associated with the expression of the following genes: *PAAF1, TBX6, ERAP1, PIP5K1C*, and *CHI3L2*. As an example, we depict plots of variant and haplotype associations with *PAAF1* (**Fig. 4B-C**), a gene linked to the regulation of proteasome function. Happler identified a candidate fine-mapped haplotype in the *PAAF1* locus spanning chr11:73935581-73910990 in GRCh38, defined by the CA allele of 11:73910990:CA:C and the TTTC allele of 11:73935581:T:TTTC. To investigate the contribution of this haplotype to the overall eQTL signal in the locus, we plotted eQTL summary statistics for the locus and conditioned on either the haplotype itself, or on one or both of its component alleles. We observed that the original signal (**Fig. 4B**, left panel) is not reduced after conditioning on the individual alleles of each haplotype (**Figs. 4B**, second and third panels). We observed an intermediate reduction when conditioning on each allele of the haplotype together (**Fig. 4B**, fourth panel). On the other hand, the signal is completely removed when conditioning on the haplotype itself (**Fig. 4B**, fifth panel). The haplotype in this locus also exhibited a stronger association than its individual alleles in GTEx (haplotype: p=2.26e-07, 11:73910990:CA:C: p=0.66, 11:73935581:T:TTTC: p=9.75e-04) (**Fig. 4C**). Overall this is consistent with the locus being best explained by the haplotype defined by these two alleles rather than by independent contributions of each variant.

**Fig 4.**
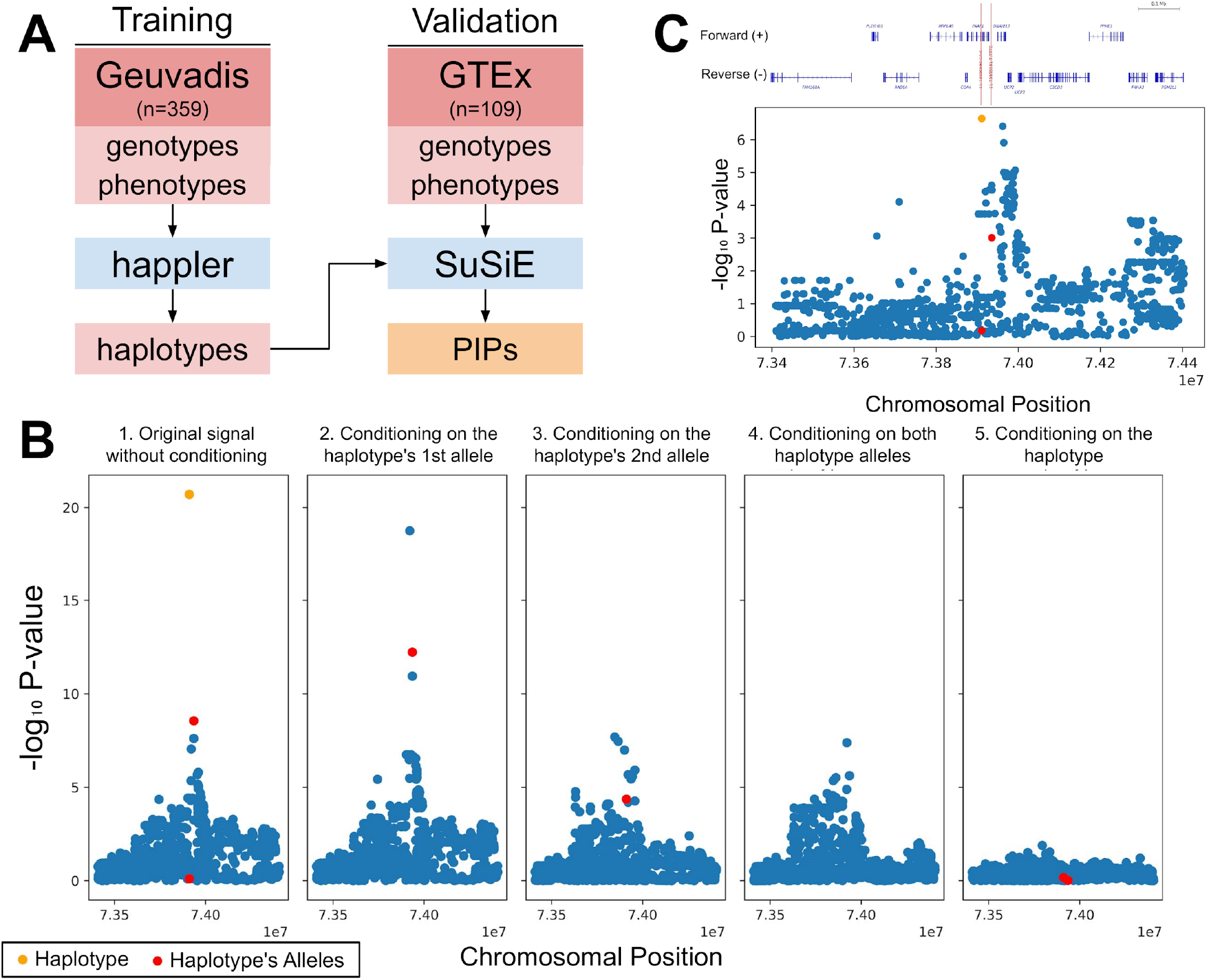
Fine-mapping eQTLs using happler. A. Schematic of Geuvadis analysis. We applied happler to 2,440 eQTL loci identified in LCL data from Geuvadis. We used European samples from Geuvadis (n=359) as a training cohort for haplotype discovery. We computed the PIPs of each haplotype by fine-mapping them alongside individual variants with SuSiE in a separate validation cohort (GTEx; n=109). **B. Summary statistics for an example locus in Geuvadis when conditioning on a haplotype vs its alleles**. The haplotype identified consists of alleles from hg38:chr11:73935581 (T/TTTC) and hg38:chr11:73910990 (CA/C) and was fine-mapped as an eQTL for the gene *PAAF1*. The original eQTL signal in Geuvadis (**B**, first panel) is not reduced compared with conditioning on each of the haplotype’s individual alleles (**B**, second and third panels) or on both alleles together (**B**, fourth panel). However, the signal is completely removed when conditioning on the candidate haplotype identified by happler (**B**, fifth panel). In each plot, blue dots show association statistics (-log10 P-values) for individual variants. Red dots show association statistics of the individual alleles of the haplotype. The orange dot represents the haplotype itself (and is plotted at the coordinate of the left-most variant in the haplotype). **C. Summary statistics for the example locus in GTEx**. The same haplotype in GTEx has a stronger association (p=2.26e-07, PIP=0.985) than any individual allele in the 1 Mbp locus.

### Haplotype tagging of rare causal variants in AoU

We set out to identify candidate haplotypes for platelet count in the AoU research cohort v8 (n=105,446). We focused on this trait since it is highly polygenic^37^, has data for a large number of samples, and also harbors known rare (MAF<0.05) and ancestry-specific variants that might be well-tagged by individual haplotypes^36^. We executed happler’s haplotype construction step in each of the 201 platelet-count GWAS loci identified by Mirmira et al.^36^ on variants with MAF>0.0001 but did not identify any candidate haplotypes using default criteria. We observed that runtime and memory usage of this step scaled linearly with the number of variants in each locus (**Supplementary Fig. 5)**.

We hypothesized that haplotypes identified by happler in the gene expression analyses above tend to tag causal variants that are rare and thus not directly included in the analysis. Due to the low sample size for the Geuvadis datasets, we had restricted analysis to variants with MAF >0.1, and thus suspect some candidate haplotypes may tag individual causal variants with MAF below this threshold. On the other hand, the scale of AoU enabled us to consider far lower allele frequencies, and thus it is possible many causal variants are directly captured and do not require haplotype analysis to identify.

We sought to investigate how the input variant MAF threshold would change the candidate haplotypes that happler produces. To assess this, we first identified high-confidence candidate causal variants with SuSiE, considering all variants with MAF>0.0001, choosing the variant with the highest PIP within each locus, and restricting to loci for which the top variant had PIP>0.9 and MAF<0.1 (**Fig 5A**). Next, we executed happler’s haplotype construction step repeatedly at each locus with increasing MAF thresholds of 0.01 to 0.1 (**Fig 5A**). The lower bound of 0.01 was chosen based on observed MAFs of high-confidence fine-mapped causal variants. This analysis yielded candidate haplotypes for seven (MAF>0.01) loci (**Fig 5C**), compared to none when using MAF>0.0001.

**Fig 5.**
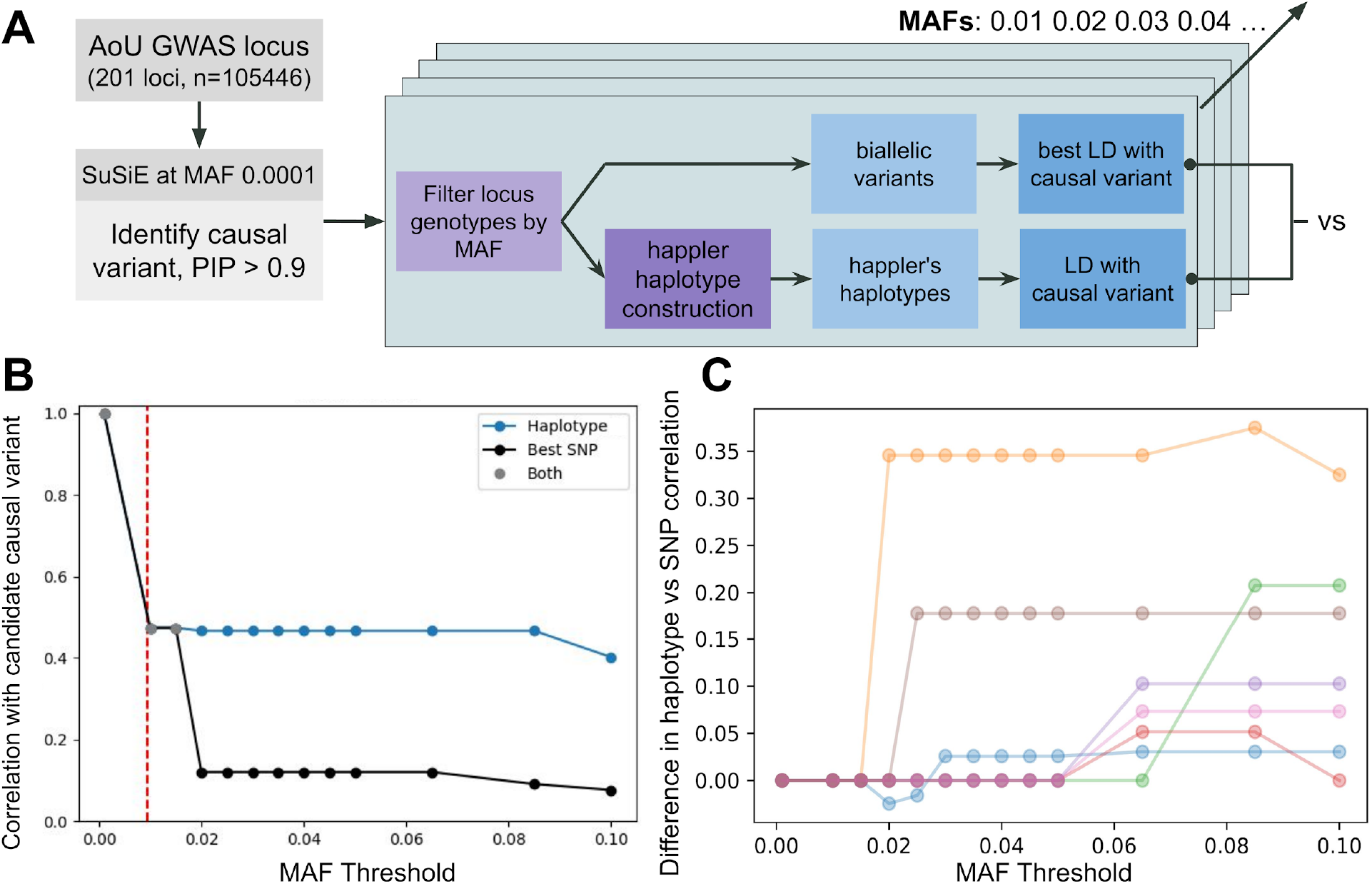
Happier identifies haplotypes tagging rare causal variants in AoU. A. Analysis flowchart. We applied happler to 201 platelet count GWAS loci in the AoU v8 dataset in a cohort of individuals self-identified as White and genetically predicted to be of European ancestry. A candidate causal variant was identified for each locus via SuSiE at an MAF>0.0001 and PIP > 0.9. At increasing MAF thresholds, we ran happler and then computed the correlation between happler’s haplotypes and the original rare candidate causal variant. We compared this to the correlation with the best biallelic variant above the MAF threshold. **B. Example locus 17:43860386-44869051**. The haplotype correlates more strongly with the rare causal variant than the best biallelic variant once the rare causal variant is filtered by MAF (dashed/red line). **C. Difference in correlation between the haplotype and the best SNP**. At high MAF thresholds, the haplotype has a stronger correlation with the rare causal variant than the best SNP. Each haplotype is assigned a different color. A total of seven haplotypes were identified by happler across the variant MAF thresholds considered. The x-axis gives the variant MAF threshold. The y-axis gives the difference in correlation with the presumed causal variant between the best haplotype and best individual variant (corresponding to the difference in the blue and black lines in panel **B**).

Across all MAF thresholds, these haplotypes generally exhibited a higher LD with the candidate causal variant than the best tagging biallelic variant, especially at MAF thresholds where the candidate causal variant itself was excluded from analysis (**Fig 5B-C**). Overall, these results suggest that the majority of loci for which happler implicates a high confidence haplotype are likely driven by rare ungenotyped variants, which may often be captured directly in large biobanks but are still largely missing from smaller scale eQTL and trait-specific GWAS cohorts (**Discussion**).

## Discussion

Here we presented happler, a novel haplotype-based fine-mapping method. Compared to existing fine-mapping strategies, happler can identify both individual variants as well as haplotypes, or combinations of alleles in *cis*, that may harbor the causal drivers underlying association signals. We benchmark happler using simulations of haplotype effects and find that in settings with sufficient sample size it can accurately identify the simulated causal haplotypes. An exception is cases where the variants of the haplotype are in strong LD, in which case the haplotype effect cannot be distinguished from the effect of the individual variants. We further demonstrate that performing traditional variant-level fine-mapping can result in inconclusive or incorrect results when the truly causal haplotype is not included in the input set of variants, and that inclusion of candidate haplotypes in fine-mapping does not introduce false positive signals as long as the fine-mapping step is performed in an independent cohort not used for haplotype discovery.. We demonstrate the utility of happler on real data by fine-mapping eQTL signals and identify multiple signals that are better explained by haplotypes rather than individual variants. Finally, we perform happler analysis of GWAS loci for platelet count in AoU and show that haplotypes identified by happler can in some cases better tag rare causal variants than any individual common variant.

The happler method faces several limitations. First, we take a greedy approach to identify candidate haplotypes, in which we start by identifying the variant with the strongest trait association and incrementally extend the haplotype by additional variants providing the strongest haplotype signal. This is a heuristic strategy to avoid exploring the entire |2^n^| search space of *n*-variant haplotypes. Our simulations show this strategy works in most cases, but this approach may not always be optimal. Second, our haplotype construction step applies lenient thresholds on the termination tests described to identify significant haplotypes, and does not account for the large number of hypothesis tests performed throughout tree building. However, we have found that we can tolerate a high false positive rate during haplotype construction, since non-causal haplotypes identified by this stage typically are assigned low PIPs during the fine-mapping stage which is performed in an independent validation cohort.

Finally, a major remaining challenge is interpretation of the haplotype effects identified. Importantly, the individual variants defining the haplotypes identified by happler are not necessarily causal themselves. In some cases, candidate haplotypes may be in strong LD with something else that is causal but not directly observed. For example, an identified haplotype could tag an unobserved rare variant or structural variant. An alternative explanation is that the the haplotype could harbor combinations of variants that exhibit *cis* interaction effects, such as combinations of protein coding variants that individually have no effect but together alter protein function. However, confidently distinguishing between these two cases remains challenging and cannot be done statistically with happler alone.

Overall, we did not find strong evidence for *cis* interaction effects for the traits considered. For the eQTL analysis, in which we included variants with MAF>0.1, we identified a total of 5 fine-mapped haplotypes across 2,440 genes considered. We anticipate some of these may tag rarer variants filtered by the stringent MAF threshold required at the low sample size (n=359 for Geuvadis). In our platelet count analysis in AoU, which had a larger sample size (n=105,446), we did not initially identify any candidate haplotypes when considering rare variants down to MAC>=20. On the other hand, we found that using higher MAF thresholds was associated with an increased probability of producing haplotypes. These haplotypes tend to be correlated with the causal variant identified by fine-mapping at a low MAF threshold, and they serve as better proxies of the causal variant than any other SNP at those thresholds. These results argue against widespread *cis* epistatic effects, at least of common variants, which we anticipate would be captured by haplotypes regardless of the MAF threshold used. Another implication of this result is that our haplotype-based approach has the most benefit in small to medium sized datasets that do not enable direct analysis of rare variants, whereas biobank-scale cohorts where rare and ultra-rare variants can be directly captured will have less benefit from a haplotype approach. Importantly, these results are based on a single trait (platelet count) and do not rule out haplotype effects at other complex traits not considered here. Ultimately, additional work is required to understand the biological mechanisms driving the haplotype associations identified from real data.

## Data availability

Geuvadis datasets and 1000GP genotypes used in this study are available from the 1000GP website [https://www.internationalgenome.org/data-portal/data-collection/geuvadis] and [https://www.internationalgenome.org/data-portal/data-collection/30x-grch38]. Analyses are based on the GRCh38 reference genome [https://storage.googleapis.com/genomics-public-data/resources/broad/hg38/v0/Homo_sapiens_assembly38.fasta]. GTEx data was obtained from dbGaP accession number phs000424.v8.p2 and from the GTEx portal website [https://www.gtexportal.org/home/downloads/adult-gtex]. This study also used data from the *All of Us* Research Program’s Controlled Tier Dataset V8, available to authorized users on the Research Workbench.

## Code availability

Happler is available at: https://github.com/gymrek-lab/happler

## Author contributions

A.M. developed happler, performed analyses, and wrote the manuscript. M.L. performed analysis of GTEx gene expression data. T.M. performed preprocessing of AoU data. U.J. helped perform analysis of Geuvadis gene expression data. J.M. provided input on design of the happler method.Y.L. performed preprocessing of GTEx and Geuvadis data. M.G. supervised the study and helped write the manuscript.

## Acknowledgements

This work was partially supported by NIH/NHGRI grant RM1HG011558 (M.G.). A.R.M. was also supported by the National Science Foundation Graduate Research Fellowship [DGE-2038238] and grants from the National Institutes of Health [T32GM008806 and T32GM139790]. The Genotype-Tissue Expression (GTEx) Project was supported by the Common Fund of the Office of the Director of the National Institutes of Health and by NCI, NHGRI, NHLBI, NIDA, NIMH, and NINDS. The data used for the analyses described in this article were obtained from the GTEx portal and dbGaP accession number phs000424.v8.p2. We gratefully acknowledge *All of Us* and UK Biobank participants for their contributions, without whom this research would not have been possible. We also thank the National Institutes of Health’s *All of Us* Research Program for making available the participant data examined in this study. We acknowledge Graham McVicker, Matteo D’Antonio, Abraham Palmer, Tiffany Amariuta, and Armin Schwartzman for helpful discussions about the happler method.

