## Supplementary for "Haplotype-based fine-mapping of variant associations for complex traits"

### Supplementary Fig. 1

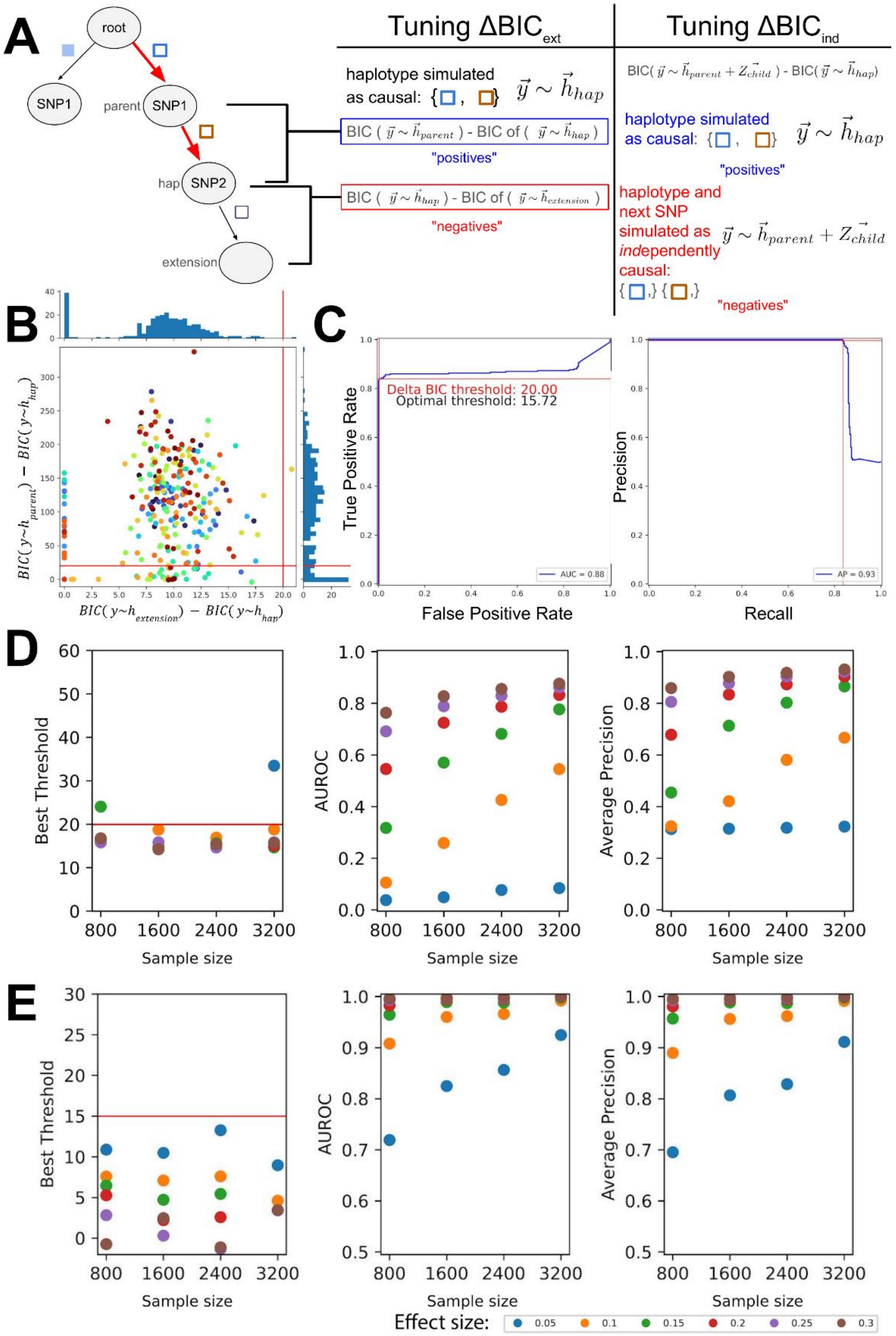

**Choosing optimal delta BIC thresholds. A.** Overview of threshold tuning. We simulated a single causal haplotype in each of 100 loci and evaluated using the $\Delta BIC_{ext}$ to classify between “positive” (correct) haplotype extensions $BIC(\, y{\sim}h_{parent}) - BIC(\, y{\sim}h_{hap})$ and “negative” (incorrect) haplotype extensions $BIC(\, y{\sim}h_{extension}) - BIC(\, y{\sim}h_{hap})$, where $h_{hap}$ is the true causal haplotype (**Methods**). We used a similar procedure for $\Delta BIC_{ind}$, where we distinguish between "positive" cases where a single haplotype (as in $y{\sim}h_{hap}$) is simulated as causal vs "negative" cases where the parent of the haplotype and the haplotype's last allele were simulated as additively causal (as in $y{\sim}h_{parent} + Z_{child}$) (**Methods**). **B. Evaluating the ability of $\Delta BIC_{ext}$ to classify correct vs. incorrect haplotype extensions.** The x-axis shows the $\Delta BIC_{ext}$ value for negative extensions and the y-axis shows the $\Delta BIC_{ext}$ value for positive extensions at each locus. We used five replicates from each locus, which are grouped by identical coloring in the scatterplot. Histograms show the distribution of values along each axis. **C. Choosing the best threshold at FDR<0.05.** We computed the receiver operator (left) and precision-recall curves (right) for using the $\Delta BIC_{ext}$ value to classify between positive and negative extension cases. For **B-C**, red lines denote the “optimal threshold” based on the minimum value of $\Delta BIC_{ext}$ that achieves a false discovery rate (FDR) <5% (corresponding here to $\Delta BIC_{ext}$=20). Data is shown for simulations with n=3200, beta=0.3. **D. Choosing a threshold for $\Delta BIC_{ext}$.** We repeated the analysis in **B-C** across a range of simulated effect sizes and sample sizes. Color denotes effect sizes. In each plot the x-axis gives the sample size. The y-axes show the optimal $\Delta BIC_{ext}$ threshold identified (left), auROC (middle), and average precision for each simulation setting. In the left panel, the red horizontal line indicates the default threshold used by happler, which provides robust results across simulation settings. **E. Choosing a threshold for $\Delta BIC_{ind}$.** We repeated the analysis in **B-D** to determine an optimal threshold for $\Delta BIC_{ind}$. Similar to **D**, panels show the optimal $\Delta BIC_{ind}$ threshold identified (left), auROC (middle), and average precision for each simulation setting.

### Supplementary Fig. 2

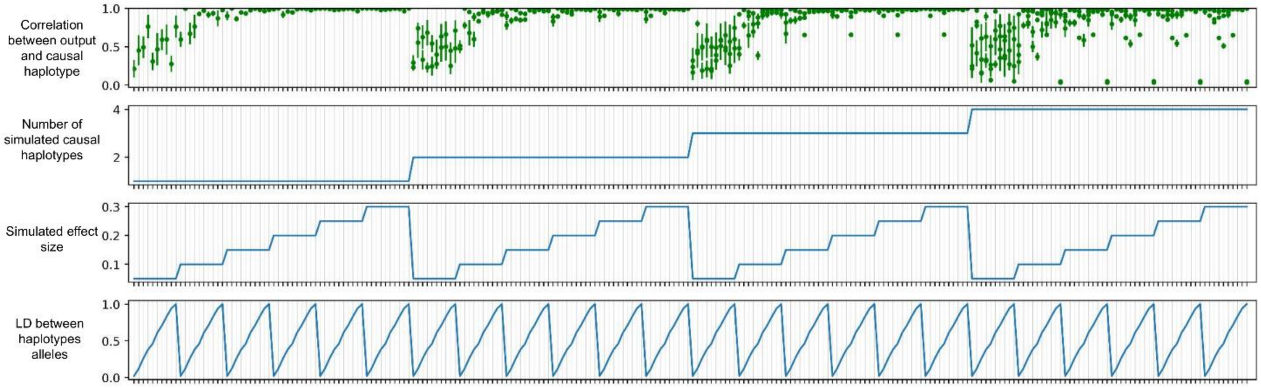


**Detailed results for evaluation of haplotype construction in the presence of simulated causal haplotypes.** As described in **Methods** we simulated phenotypes based on a single causal haplotype and performed haplotype construction using happler on genotypes within the 1Mb fine-mapping region. Each x-axis value represents a separate simulation. The bottom three panels show simulation settings. From bottom to top, panels show: (1) LD between the alleles of the causal haplotype, (2) Effect size, and (3) the number of simulated causal haplotypes. The top panel shows the LD of each output haplotype with the best matched causal haplotype. Error bars show the mean +/- 1 std. error across 10 simulations. Data is shown for a sample size of 3200.

### Supplementary Fig. 3

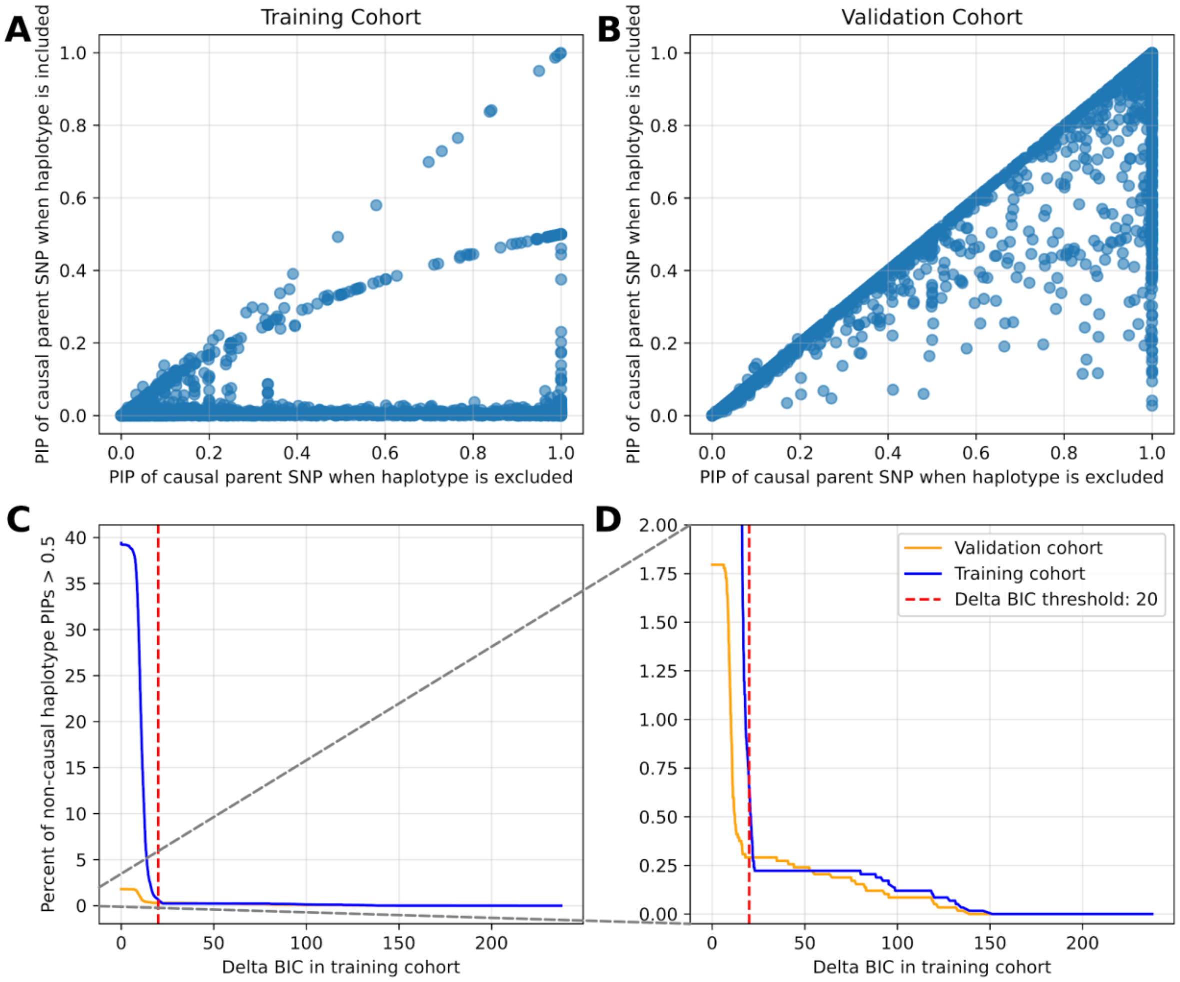


**Evaluating the use of the same cohort used for haplotype discovery (training cohort) vs. an independent validation cohort for the fine-mapping stage of happler. A** and **B** summarize fine-mapping in the training (**A**) and validation (**B**) cohorts. We simulated a single causal SNP and ran happler's *haplotype construction* in a training cohort to identify the best extension of the SNP, generating a single (false positive) haplotype. Next, we ran SuSiE in each cohort with the incorrect haplotype either included or excluded in the set of input variants. Plots show the PIP of the causal SNP when the false positive haplotype is not (x-axis) vs. is (y-axis) included as input. PIPs of the true causal SNP are greatly reduced when performing fine-mapping in the training cohort (left) but not the validation cohort (right), demonstrating the utility of using an independent cohort for fine-mapping to avoid over-fitting. Each dot represents a single simulation round with the same loci, effect sizes, and sample sizes as Supp Fig 1. **C. False positive haplotypes identified at different $\Delta BIC_{ext}$ values.** The x-axis gives the $\Delta BIC_{ext}$ threshold. The y-axis shows the percentage of false positive haplotypes with $\Delta BIC_{ext}$ values above each threshold that were fine-mapped with PIP>0.5 in the validation (orange) or training (blue) cohorts. The red vertical line shows the default $\Delta BIC_{ext}$ threshold of 20 used by happler. Panel **D** is the same as **C** but zoomed in to show y-axis detail from 0-2. The rates of false positive fine-mapped haplotypes are far lower when fine-mapping is performed in the validation cohort.

### Supplementary Fig. 4

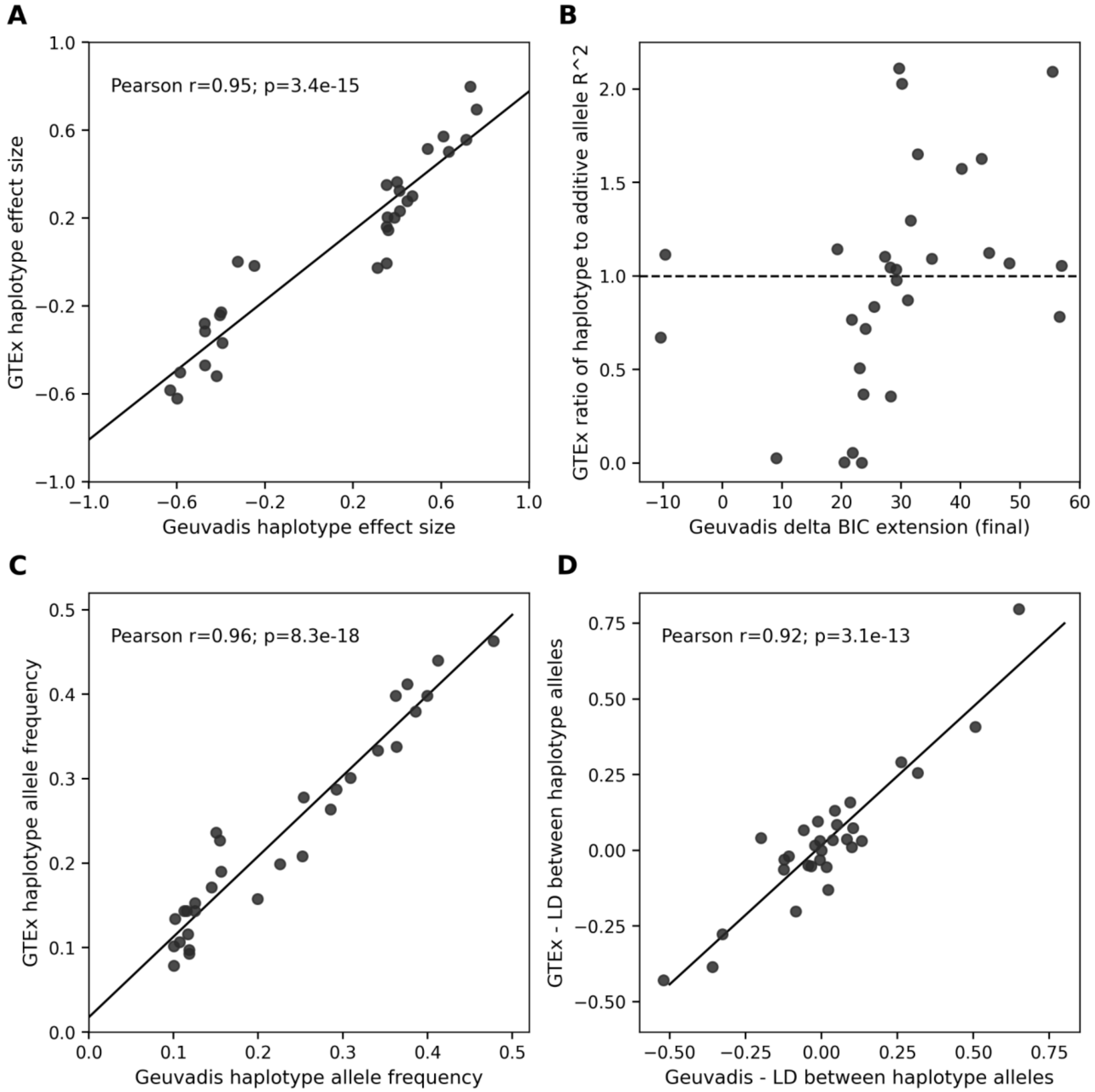


**Replication of Geuvadis haplotype associations in GTEx. A. Comparison of haplotype effect sizes**. The x-axis gives effect sizes estimated in Geuvadis and the y-axis gives those estimated in GTEx. **B. Replication of haplotype explanatory power in GTEx as a function of the haplotype's $\Delta BIC_{ext,final}$ score in the Geuvadis training cohort.** The y-axis, computed in GTEx, is the ratio of the $R^2$ of the haplotype model relative to the $R^2$ of a model where the two constituent alleles of the haplotype are additively associated. The horizontal dashed line denotes y=1. **C. Comparison of haplotype allele frequencies between Geuvadis and GTEx cohorts.** The x-axis and y-axis show the haplotype allele frequency in Geuvadis and GTEx, respectively. **D. Comparison of linkage disequilibrium (computed as the unsquared pearson correlation**

**coefficient) between constituent haplotype alleles in Geuvadis versus GTEx.** The x-axis and y-axis show the LD values in Geuvadis and GTEx, respectively for each haplotype. Pearson correlation coefficients ($r$) and corresponding two-sided $p$-values are reported for panels A, C, and D. In all cases, the solid black line indicates the ordinary least squares (OLS) regression line of best fit.

### Supplementary Fig. 5

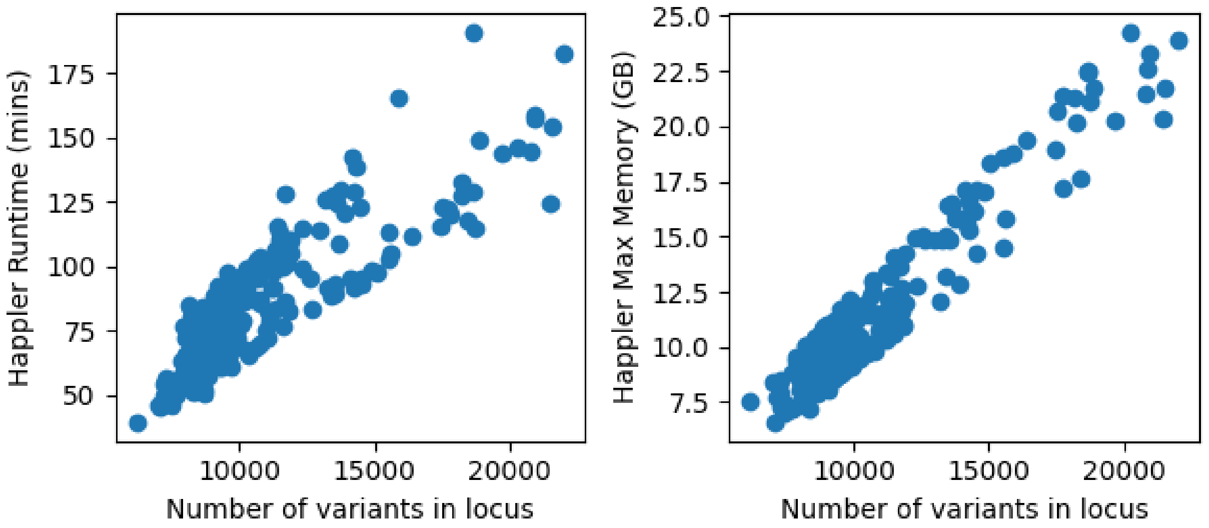


**Runtime and memory usage of the *haplotype construction* stage in AoU.** Runtime and memory were assessed for each of 201 platelet-count associated loci in AoU v8 (n=105,446). Runtime and memory usage scale linearly with the number of variants in the fine-mapping locus. The memory usage can be reduced, while trading-off a subsequent increase in runtime, by increasing the value provided to --chunk-size, which controls how many variants are processed by happler simultaneously.
